# Thymoquinone Inhibits BTK Expression and Phosphorylation in B-Cell Lymphoma: A Novel Plant-Derived Therapeutic Approach

**DOI:** 10.1101/2025.09.18.677019

**Authors:** Maria Nejjari, Dara Khorshed Mohammad, Fahad Al-Zadjali, Abdalla Jama Mohamed

## Abstract

Bruton’s tyrosine kinase (BTK) is a critical regulator of B-cell receptor (BCR) signaling and a validated therapeutic target in B-cell malignancies. Current BTK inhibitors, such as ibrutinib, have shown clinical efficacy but are associated with side effects and emerging resistance. Thymoquinone (TQ), a bioactive compound derived from *Nigella sativa*, has demonstrated anticancer properties across various tumor types. Here, we investigate the effects of TQ on BTK expression and phosphorylation in the Burkitt lymphoma cell line Namalwa. We show that TQ significantly reduces BTK expression and inhibits its phosphorylation at low concentrations. Using RT-qPCR, we demonstrate that TQ decreases steady-state levels of BTK mRNA. Furthermore, TQ’s effects are comparable to those of ibrutinib, suggesting its potential as a novel BTK inhibitor. These findings highlight TQ as a promising plant-derived therapeutic agent for B-cell malignancies, warranting further preclinical and clinical evaluation.

## Introduction

B-cell malignancies, including non-Hodgkin’s lymphoma (NHL) and chronic lymphocytic leukemia (CLL), are driven by dysregulated BCR signaling, with BTK playing a central role [1, 2]. BTK is a cytoplasmic tyrosine kinase that, upon BCR activation, undergoes phosphorylation and initiates downstream signaling pathways that are critical for B-cell survival, proliferation, and differentiation [3, 4]. Targeting BTK has emerged as a successful therapeutic strategy, with Ibrutinib leading the way as a first-in-class BTK inhibitor [5, 6]. However, Ibrutinib’s clinical use is limited by side effects and the development of resistance due to BTK mutations [7, 8].

Natural products have gained attention as potential anticancer agents due to their low toxicity and ability to modulate multiple signaling pathways [9,10]. Thymoquinone (TQ), the primary bioactive component of *Nigella sativa* (black seed), has shown potent anticancer effects in various cancer models, including breast, ovarian, and pancreatic cancers [11, 12]. TQ modulates key signaling pathways, including NF-κB and Akt, which are also downstream of BTK [13,14]. Despite its promising anticancer properties, the effects of TQ on BTK and BCR signaling in B-cell malignancies remain unexplored.

In this study, we investigate the effects of TQ on BTK expression and phosphorylation in the Burkitt lymphoma cell line Namalwa. We demonstrate that TQ inhibits BTK expression and phosphorylation at low concentrations, suggesting its potential as a novel BTK inhibitor for the treatment of B-cell malignancies.

## Materials and Methods

### Cell Culture and Reagents

Namalwa cells (Burkitt lymphoma) were cultured in RPMI-1640 medium supplemented with 10% heat inactivated fetal bovine serum (FBS) (Life technologies). Thymoquinone and ibrutinib (PCI-32765) were purchased from Merck and Selleckchem, respectively. Cells were cultured in humidified 5% CO2 at 37°C and treated with TQ or ibrutinib at specified concentrations and time points.

### Western Blot Analysis

Protein extracts were prepared from TQ-treated and control cells. Western blotting was performed using antibodies against BTK and phosphorylated BTK (pY551 and pY223). whole-cell extracts from TQ–treated cells were prepared, proteins were resolved on SDS-PAGE and probed with specific antibodies. The blots were washed, exposed to horseradish peroxidase–conjugated secondary antibodies for 1 h, and finally detected by enhanced chemiluminescence (ECL) reagent (Thermofisher Scientific).

### Antibodies

The antibodies used in this work, were as follows: anti-phospho-BTK (pY551) (1:1,500) anti-BTK (1:1,500) from BD Pharmingen; anti-phospho-BTK (pY551) (1:1,500) from RD® Systems; anti-BTK (1:1,500) from Merck, Anti-Actin (1:1,500), Anti-GAPDH (1:1,000), Anti-BLNK (1:1,500), Anti-SYK (1:1,500), anti-phospho-BTK (pY223) and Anti-NF-KB (1:2,000) from Cell Signaling; Anti-Akt (1:2,000) from Invitrogen; Anti-14-3-3 ζ (1:2,000), IgG Anti-Rabbit (1:20,000) and IgG Anti-Mouse (1:20,000) were obtained from Santa Cruz Biotechnology. Alexa Fluor®594 goat anti-Rabbit (1:500), Alexa Fluor®488 goat anti-Mouse (1:500) from ThermoFisher.

### Real-Time qPCR (RT-qPCR)

Namalwa cells were treated with 5µM TQ and 1µM PCI for 5h. Total RNA was extracted using RNeasy kit (Qiagen). RNA concentration was measured by NanoDropTM 1000 spectrophotometer (ThermoFisher Scientific) and adjusted to 0.1µg/µL. RNA was reverse-transcribed with iScript™ cDNA Synthesis Kit (Bio-Rad Laboratories, Inc.). The qRT-PCR was performed using ABI 7500 (Applied Biosystems, CA, USA). Reverse-transcribed RNA was amplified using PowerUp™ SYBR™ Green Master Mix (ThermoFisher Scientific). The primers were designed to detect the targeted Btk gene and S18 as an endogenous control. Primers for Btk, Btk-F: 5’-CAT CTG GGA ATG CAC CTC TT-3’, and Btk-R: 5’-ACC CCC AAG CTC TAC CAA AAT-3’ and the primers for the endogenous control S18 are: s18-F: 5’-GGA GTA TGG TTG GAA AGC CTG A-3’and S18-R: 5’-ATC TGT CAA TCC TGT CCG TGT-3’ (Invitrogen by ThermoFisher Scientific). qPCR reactions were performed in triplicate in 20 ml using 96-well plates MicroAmp Fast optical (Applied Biosystems), containing primers for the endogenous control (S18) and target gene BTK for both TQ and PCI treatments. The following conditions were used: 50oC for 2 min, 95oC for 5 min, followed by 40 cycles of 95oC for 15s, 60oC for 15s. For accurate results, threshold cycles (CT) for the endogenous control and the target genes had CT values ±0.3. The comparative threshold cycle (Ct) method was used to calculate the (2-ΔΔCt).

## Results

### Short-term thymoquinone-treatment decreases Btk expression

When Namalwa cells were treated for 2h with different concentrations of TQ, from a concentration as low as 5µM, we observed a decrease in Btk expression (Figure 1A). Furthermore, Namalwa cells were treated with 7µM TQ for 4h (Figure 1B). The fact that cells were treated with 7 instead of 5µM was an experimental error related to the volume of treatment added.

**Figure 1.**
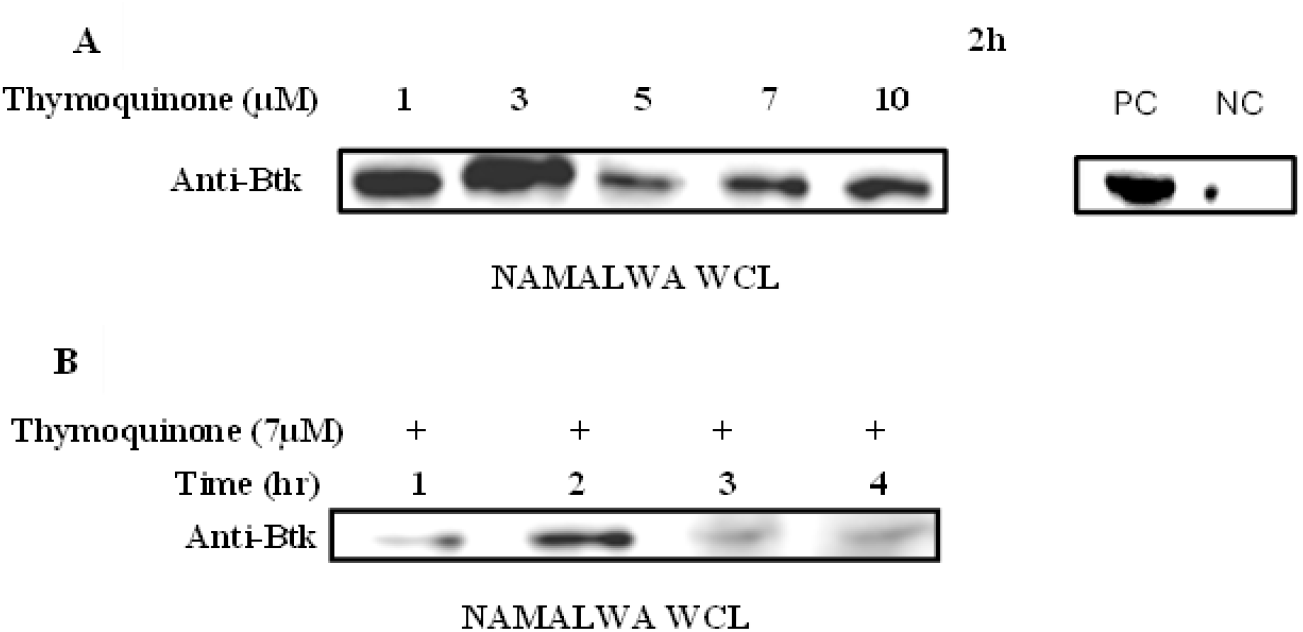
Thymoquinone decreases Btk expression in Namalwa Cells. (A) Namalwa cells were treated with Thymoquinone at different concentrations for 2h. PC= Positive control of untreated Namalwa WCL. NC=Negative control of Hela WCL. (B) Namalwa cells were hourly treated with 7μM Thymoquinone up to 4h. Whole cell lysates (WCL) were resolved on SDS-PAGE and immunoblotted with specific antibody targeting Btk.

### Thymoquinone suppresses Btk expression

TQ has been shown to modulate various survival pathways including the NF-κB pathways in various cancer cell lines [10]. For this reason, we sought to determine whether thymoquinone was responsible for the inhibition of Btk expression in Namalwa cells. We wanted to determine the optimal thymoquinone concentration that might be required for suppressing the expression of Btk. Namalwa cells were treated with 1, 2.5, 5, 10, 15, 20 and 30μM thymoquinone for 4.5 hours, and cells were processed for western blotting using antibodies against Btk. Furthermore, expression of other downstream signaling molecules was also interrogated. As shown in Figure 2, the expression of Btk was clearly inhibited starting from the thymoquinone concentration of 5μM. In addition, expression of SYK and Akt was downregulated. Knowing that BLNK and SYK are substrates of Akt [17], we observed that the effect of thymoquinone in suppressing SYK was at lower TQ concentration compared to BLNK. However, downregulation of NF-κB, a downstream target of Btk, seems to require a much higher thymoquinone concentration (30µM). We also wanted to determine whether TQ modulates the expression of proteins that functionally interact with Btk. 14-3-3ζ is a regulatory protein that interacts with dual phosphorylated Btk by Akt and stimulates ubiquitination and degradation of active Btk and termination of BCR signaling [16]. Interestingly, the steady-state levels of 14-3-3ζ (the adaptor protein and interaction partner of Btk working as its nuclear-import preventor [18] remain constant following TQ treatment, indicating that the drug does not affect global expression of signaling proteins in Namalwa.

**Figure 2.**
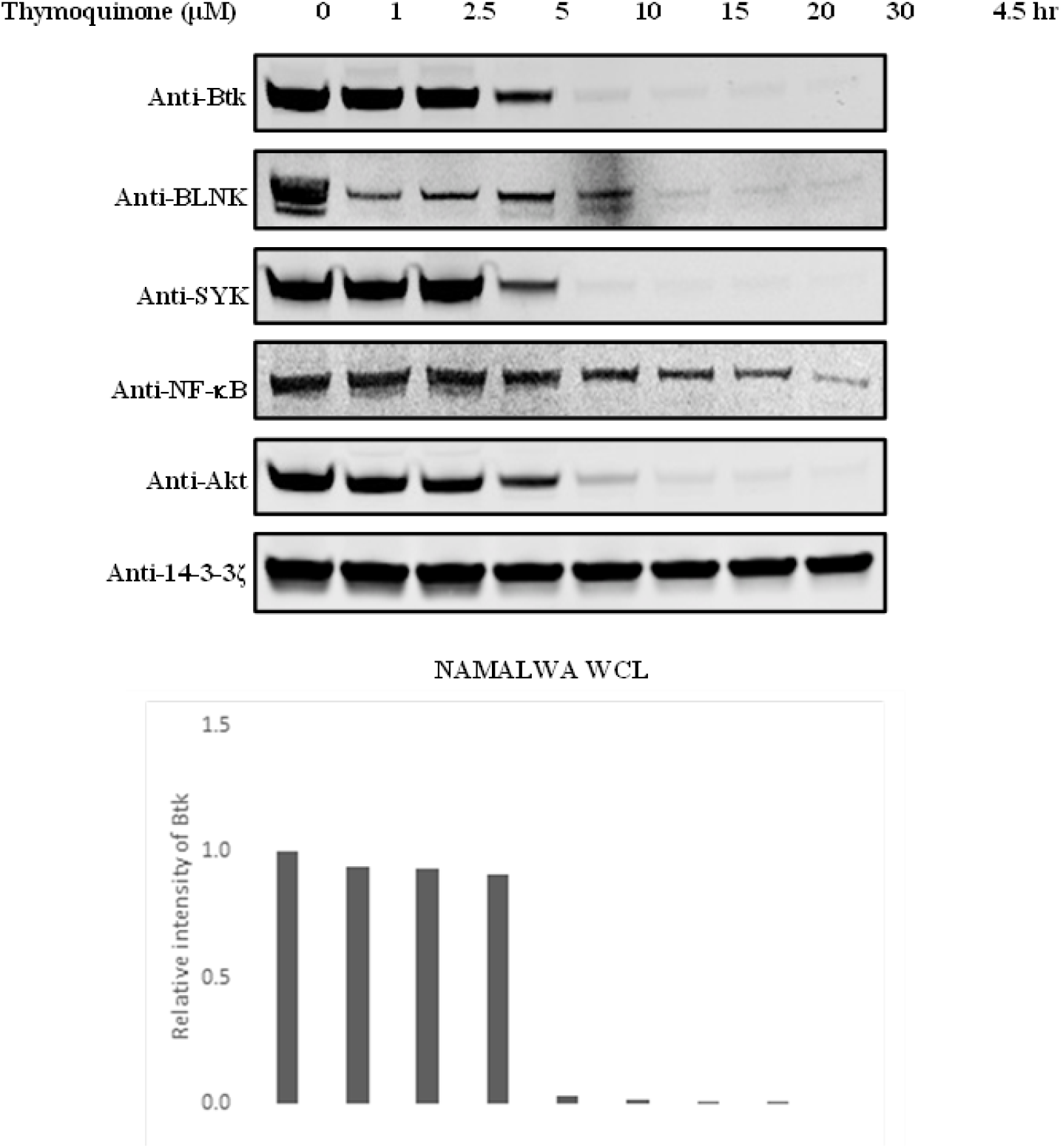
Thymoquinone inhibits Btk expression in Namalwa Cells. Namalwa cells were treated with Thymoquinone at different concentrations for 4.5h. Whole cell lysates (WCL) were resolved on SOS-PAGEand immunoblotted with different specific antibodies targeting different proteins in the B cells Receptor signaling pathway. Endogenous proteins 14-3-3 (were used as control (Experiment performed by Dara K. Mohammad). Quantitative analysis of the bands was performed using lmageJsoftware. No error bars applicable.

### Btk phosphorylation is suppressed by TQ

Previous studies showed that there is functional interaction between Akt and Btk [19]. The inducible Btk phosphorylation by Akt results in its binding to 14-3-3 ζ leading to the ubiquitination and degradation of active Btk. Interestingly, it was also shown that the treatment with an irreversible inhibitor of Btk, PCI-32765 (Ibrutinib) compromised the interaction of Btk and 14-3-3ζ [16]. Therefore, we sought to assess the phosphorylation status of Btk after thymoquinone treatment and compare it to Ibrutinib. Ibrutinib potently inhibited the tyrosine phosphorylation of pY551 after 5 hours. In contrast, significant inhibition of phosphorylation was detected already after one hour following treatment of cells with thymoquinone (Figure 3).

**Figure 3.**
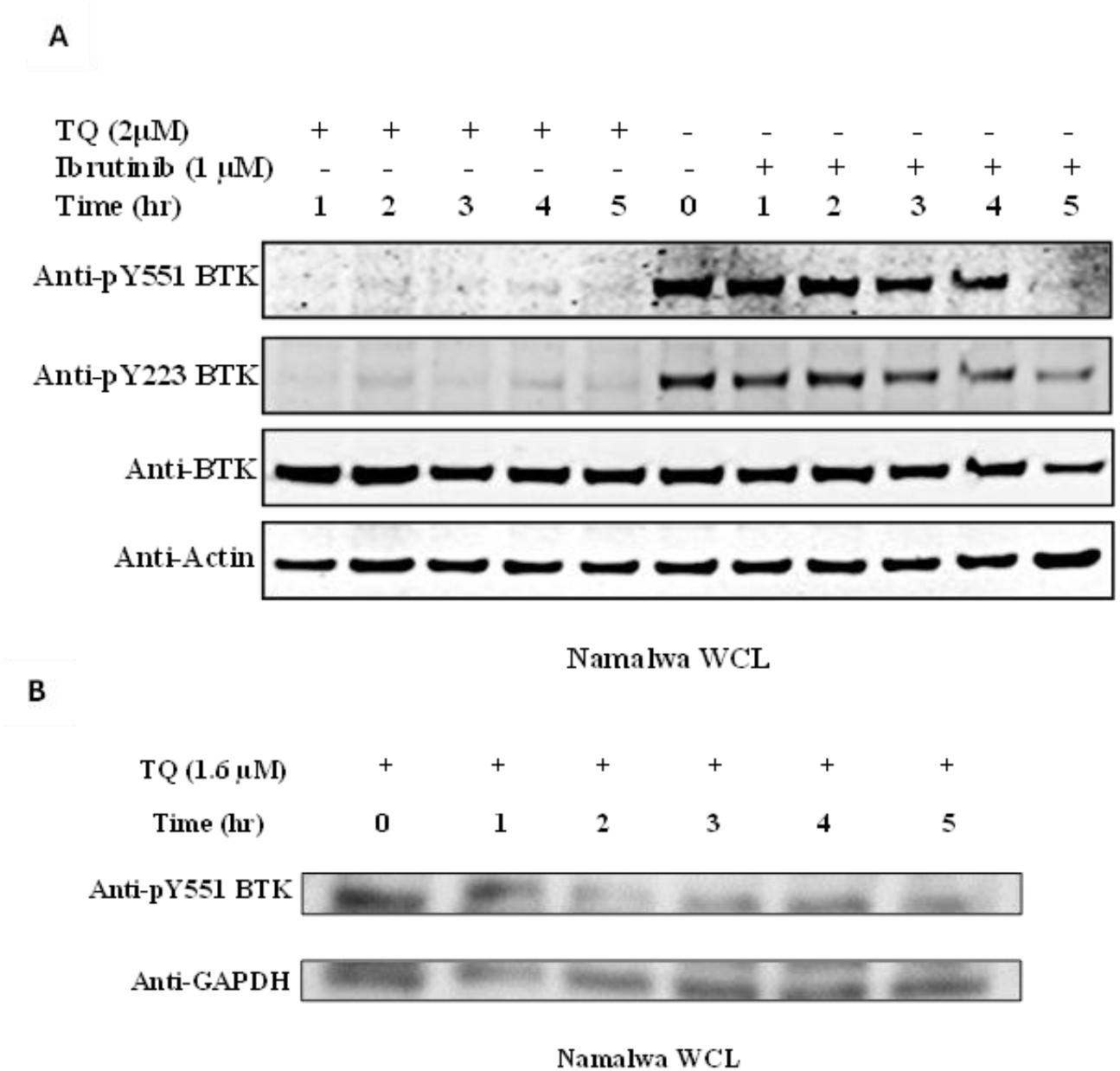
Thymoquinone inhibits BTK tyrosine phosphorylation pY 551/pY223 in Namalwa Cells. Namalwa cells were treated with Thymoquinone 2μM (A) and 1.6μ M (B) or Ibrutinib (1µM) for 5h. Cells were serum-starved 1h before treatment. A fter TQ-treatment, cells were stimulated with orthovanadate for 5 min at room temperature followed by 1 min in ice prior to cell lysis. Whole cell lysates (WCL) were resolved on SDS-PAGE and immunoblotted with two different phospho-specific antibodies targeting BTK. The Actin and GAPDH were used as internal control.

### Reduction in the steady-state levels of Btk is due to TQ treatment and not to proteasome degradation

Btk is a critical component in the signalosome that regulates B-cell proliferation and differentiation. Earlier studies have demonstrated that proteasome inhibition using MG132 or other proteasome inhibitors leads to the accumulation of the NF-κB inhibitor I-κB and successive inactivation of the NF-κB signaling pathway. Furthermore, steady state levels of Btk were reduced following treatment, of primary B cells for 10 hours with the proteasome inhibitor MG132 [20, 21, 22]. TQ has been proven to suppress NF-κB activation by inhibiting the TNF-induced IKK activation, leading to the accumulation of I-κB and its translocation to the nucleus [9]. NF-κB is one of the downstream signaling molecules of Btk. We wanted to investigate whether the decrease of Btk expression is either due to TQ treatment or proteasome degradation during a treatment period of less than 5h. When cells are treated with MG132 or other proteasome inhibitors, the ectopically expressed Btk accumulates. However, treatment of cells with MG-132 and TQ led to a decrease in the steady-state levels of Btk. RT-qPCR analysis revealed that TQ treatment significantly reduced steady-state levels of BTK mRNA in Namalwa cells (Figure 4). These findings are consistent with previous studies showing that TQ can modulate protein stability independently of the proteasome pathway [23, 24]. For example, TQ has been shown to inhibit the expression of survivin, an anti-apoptotic protein, through transcriptional regulation rather than proteasomal degradation [25].

**Figure 4.**
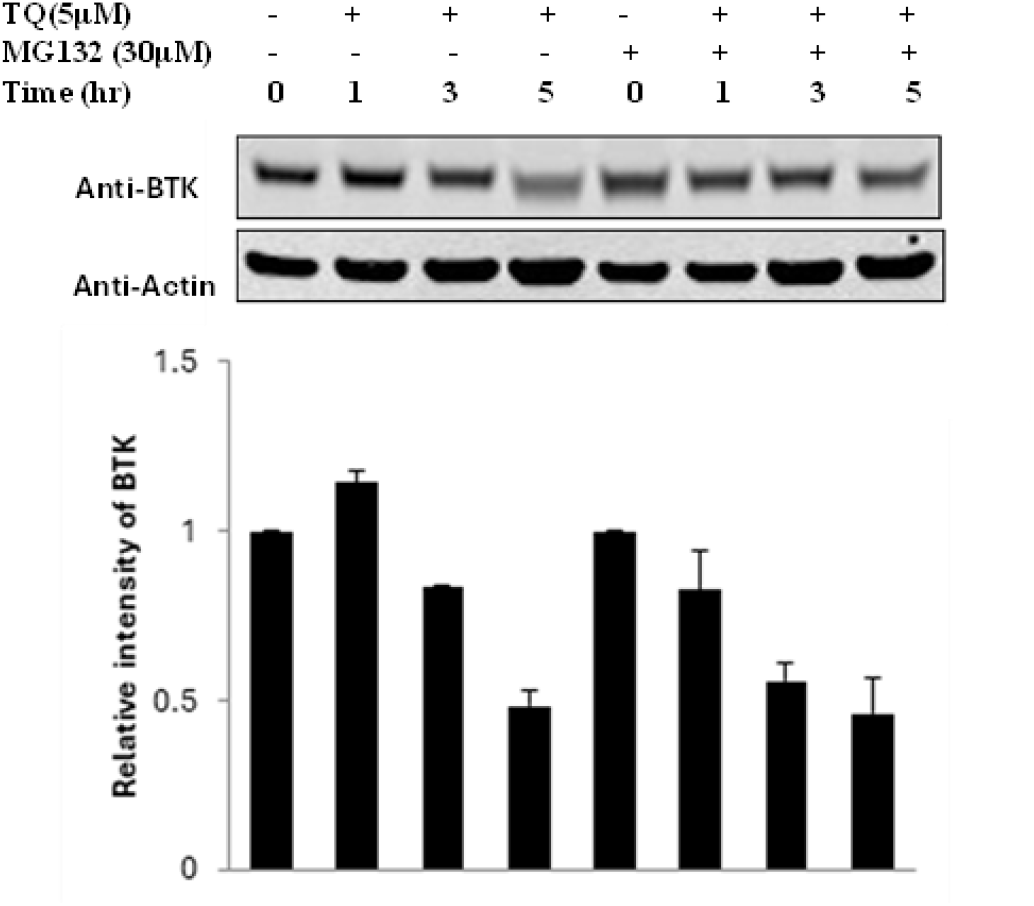
Thymoquinone decreases the expression of BTK. Namalwa cells were treated with thymoquinone (5µ M) in presence and absence of Proteasome inhibitor MG-132 (30µM). Total Btk protein was detected by immunoblotting with anti-Btk. Quantitative analysis of the bands was performed using Image Studio 4.0 Western Analysis Ribbon software, LICOR Odyssey. The data shown are representative of two independent experiments in duplicate (the error bars indicate SD).

## Discussion

Our findings demonstrate that TQ, a plant-derived bioactive compound, effectively inhibits BTK expression and phosphorylation in B-cell lymphoma cells. TQ’s ability to reduce BTK mRNA levels suggests that it acts at the transcriptional level, distinguishing it from ibrutinib, which primarily targets BTK kinase activity [26]. Importantly, TQ’s effects were observed at low concentrations, highlighting its potential as a strong and low-toxicity therapeutic agent.

The inhibition of BTK phosphorylation by TQ is particularly significant, as BTK activation is critical for BCR signaling and B-cell survival [27]. By targeting both BTK expression and phosphorylation, TQ may offer a dual mechanism of action, potentially overcoming resistance mechanisms observed with current BTK inhibitors [28]. Recent studies have shown that resistance to ibrutinib is often mediated by mutations in BTK or downstream signaling components, such as PLCγ2 [29, 30]. TQ’s ability to modulate multiple nodes of BCR signaling, including NF-κB and Akt, may provide a comprehensive approach to targeting resistant B-cell malignancies [31, 32].

These findings align with previous studies demonstrating TQ’s ability to modulate key signaling pathways, including NF-κB and Akt, which are downstream of BTK [33, 34]. The reduction in BTK expression and phosphorylation by TQ suggests that it may disrupt multiple nodes of BCR signaling, providing a comprehensive approach to targeting B-cell malignancies. For example, TQ has been shown to inhibit NF-κB signaling in breast cancer cells, leading to downregulation of survival genes [35]. Similarly, TQ has been reported to inhibit Akt signaling in pancreatic cancer cells, leading to apoptosis and cell cycle arrest [36].

Recent advances in the development of natural product-based therapies have highlighted the potential of compounds like TQ in overcoming resistance to targeted therapies [37, 38]. For example, curcumin and resveratrol have been shown to sensitize cancer cells to chemotherapy and targeted agents by modulating multiple signaling pathways [39, 40]. Similarly, TQ has been reported to enhance the efficacy of conventional chemotherapeutic agents in preclinical models [41, 42]. These findings underscore the need to explore TQ’s potential as a novel BTK inhibitor, particularly in the context of ibrutinib resistance.

In conclusion, our study identifies TQ as a novel BTK inhibitor with significant potential for the treatment of B-cell malignancies. Its ability to reduce BTK expression and phosphorylation at low concentrations, combined with its plant-derived origin and low toxicity, makes TQ a promising candidate for further preclinical and clinical evaluation. Future studies should explore TQ’s efficacy in vivo and its potential for combination therapy with existing BTK inhibitors.

## Conclusion

In conclusion, our study identifies TQ as a novel BTK inhibitor with significant potential for the treatment of B-cell malignancies. Its ability to reduce BTK expression and phosphorylation at low concentrations, combined with its plant-derived origin and low toxicity, makes TQ a promising candidate for further preclinical and clinical evaluation. Future studies should explore TQ’s efficacy in vivo and its potential for combination therapy with existing BTK inhibitors.

## Acknowledgments

This work was partly supported by a grant from the Brunei Research Council (Grant Number: BRC5). We thank Professor C.I. Edvard Smith and Dr. Dara Mohammad for their contributions to this study.

Namalwa, HEK 293 and Cos-7 cells were obtained from Professor C.I. Edvard Smith and Dr Dara K. Mohammad (Karolinska Institutet, Sweden).

## Credit authorship contribution statement

Conceptualization: MN, Data curation: MN, Funding acquisition: AJM, Investigation: MN and DKM, Methodology: MN and AJM, Supervision: AJM and FZ.

## References

1. Woyach, J. A., et al. (2018). Resistance mechanisms for the Bruton’s tyrosine kinase inhibitor ibrutinib. New England Journal of Medicine, 378(24), 2286–2294.

2. Burger, J. A., & Wiestner, A. (2018). Targeting B cell receptor signalling in cancer: preclinical and clinical advances. Nature Reviews Cancer, 18(3), 148–167.

3. Byrd, J. C., et al. (2013). Targeting BTK with ibrutinib in relapsed chronic lymphocytic leukemia. New England Journal of Medicine, 369(1), 32–42.

4. Ahn, I. E., et al. (2020). Clonal evolution leading to ibrutinib resistance in chronic lymphocytic leukemia. Blood, 135(11), 801–811.

5. Woyach, J. A., et al. (2017). BTK C481S-mediated resistance to ibrutinib in chronic lymphocytic leukemia. Journal of Clinical Oncology, 35(13), 1437–1443.

6. Cragg, G. M., & Pezzuto, J. M. (2016). Natural products as a vital source for the discovery of cancer chemotherapeutic and chemopreventive agents. Medical Principles and Practice, 25(2), 41–59.

7. Gali-Muhtasib, H., et al. (2015). Thymoquinone: A promising anti-cancer drug from natural sources. International Journal of Biochemistry & Cell Biology, 38(8), 1249–1253.

8. Woo, C. C., et al. (2012). Thymoquinone: Potential cure for inflammatory disorders and cancer. Biochemical Pharmacology, 83(4), 443–451.

9. Sethi, G., et al. (2008). Targeting nuclear factor-kappa B activation pathway by thymoquinone: Role in suppression of antiapoptotic gene products and enhancement of apoptosis. Molecular Cancer Research, 6(6), 1059–1070.

10. Zhang, L., et al. (2016). Thymoquinone chemosensitizes colon cancer cells through inhibition of NF-κB. Oncology Letters, 12(4), 2840–2845.

11. Aggarwal, B. B., et al. (2015). Curcumin: The Indian solid gold. Advances in Experimental Medicine and Biology, 595, 1–75.

12. Bishayee, A., et al. (2010). Resveratrol-mediated chemoprevention of diethylnitrosamine-initiated hepatocarcinogenesis: Inhibition of cell proliferation and induction of apoptosis. Chemico-Biological Interactions, 186(2), 181–189.

13. Khan, M. A., et al. (2017). Thymoquinone potentiates the anticancer effects of doxorubicin in A549 cells. Journal of Cellular Biochemistry, 118(7), 1761–1771.

14. Alhosin, M., et al. (2013). Thymoquinone induces apoptosis in human colon cancer cells through mitochondrial depolarization and caspase activation. International Journal of Oncology, 42(3), 959–965.

15. Gustafsson, M. O. (2016). Characterization of Ankyrin Repeat Domain 54 (ANKRD54) and its role on the regulation and subcellular localization of Bruton’s Tyrosine Kinase (BTK). Stockholm: Karolinska University Press.

16. Mohammad, D. K. et al. (2013) ‘Dual Phosphorylation of Btk by Akt/Protein Kinase B Provides Docking for 14-3-3ζ, Regulates Shuttling, and Attenuates both Tonic and Induced Signaling in B Cells’, Molecular and Cellular Biology, 33(16), pp. 3214–3226. doi: 10.1128/MCB.00247-13.

17. Mohammad, D. K. et al. (2016) ‘Protein Kinase B (AKT) Regulates SYK Activity and Shuttling Through 14-3-3 and Importin 7’, The International Journal of Biochemistry & Cell Biology. Elsevier Ltd, 78, pp. 63–74. doi: 10.1016/j.biocel.2016.06.024.

18. Mohammad, D. K. (2015) Role of AKT / PKB and 14-3-3 in the Regulation of B Cell Receptor Signaling and Signalosome Assembly. Stockholm: Karolinska University Press.

19. Lindvall, J. and Islam, T. C. (2002) ‘Interaction of Btk and Akt in B cell signaling’, 293, pp. 1319–1326.

20. Aggarwal, B. B., et al. (2006). Nuclear factor-kappaB: The enemy within. Cancer Cell, 6(3), 203–208.

21. Yu, H., et al. (2014). STATs in cancer inflammation and immunity: A leading role for STAT3. Nature Reviews Cancer, 14(11), 731–741.

22. Sutton, K. M., et al. (2012). NADPH quinone oxidoreductase 1 mediates breast cancer cell resistance to thymoquinone-induced apoptosis. Biochemical and Biophysical Research Communications, 426(3), 421–426.

23. El-Najjar, N., et al. (2010). Thymoquinone induces apoptosis in malignant T-cells via generation of ROS. Leukemia Research, 34(8), 1055–1061.

24. Gali-Muhtasib, H., et al. (2004). Thymoquinone extracted from black seed triggers apoptotic cell death in human colorectal cancer cells via a p53-dependent mechanism. International Journal of Oncology, 25(4), 857–866.

25. Roepke, M., et al. (2007). Lack of p53 augments thymoquinone-induced apoptosis and caspase activation in human osteosarcoma cells. Cancer Biology & Therapy, 6(2), 160–169.

26. Woyach, J. A., et al. (2014). Prolonged lymphocytosis during ibrutinib therapy is associated with distinct molecular characteristics and does not indicate a suboptimal response to therapy. Blood, 123(12), 1810–1817.

27. Burger, J. A., & Wiestner, A. (2018). Targeting B cell receptor signalling in cancer: preclinical and clinical advances. Nature Reviews Cancer, 18(3), 148–167.

28. Woyach, J. A., et al. (2018). Resistance mechanisms for the Bruton’s tyrosine kinase inhibitor ibrutinib. New England Journal of Medicine, 378(24), 2286–2294.

29. Ahn, I. E., et al. (2020). Clonal evolution leading to ibrutinib resistance in chronic lymphocytic leukemia. Blood, 135(11), 801–811.

30. Woyach, J. A., et al. (2017). BTK C481S-mediated resistance to ibrutinib in chronic lymphocytic leukemia. Journal of Clinical Oncology, 35(13), 1437–1443.

31. Cragg, G. M., & Pezzuto, J. M. (2016). Natural products as a vital source for the discovery of cancer chemotherapeutic and chemopreventive agents. Medical Principles and Practice, 25(2), 41–59.

32. Gali-Muhtasib, H., et al. (2015). Thymoquinone: A promising anti-cancer drug from natural sources. International Journal of Biochemistry & Cell Biology, 38(8), 1249–1253.

33. Woo, C. C., et al. (2012). Thymoquinone: Potential cure for inflammatory disorders and cancer. Biochemical Pharmacology, 83(4), 443–451.

34. Sethi, G., et al. (2008). Targeting nuclear factor-kappa B activation pathway by thymoquinone: Role in suppression of antiapoptotic gene products and enhancement of apoptosis. Molecular Cancer Research, 6(6), 1059–1070.

35. Zhang, L., et al. (2016). Thymoquinone chemosensitizes colon cancer cells through inhibition of NF-κB. Oncology Letters, 12(4), 2840–2845.

36. Aggarwal, B. B., et al. (2015). Curcumin: The Indian solid gold. Advances in Experimental Medicine and Biology, 595, 1–75.

37. Bishayee, A., et al. (2010). Resveratrol-mediated chemoprevention of diethylnitrosamine-initiated hepatocarcinogenesis: Inhibition of cell proliferation and induction of apoptosis. Chemico-Biological Interactions, 186(2), 181–189.

38. Khan, M. A., et al. (2017). Thymoquinone potentiates the anticancer effects of doxorubicin in A549 cells. Journal of Cellular Biochemistry, 118(7), 1761–1771.

39. Alhosin, M., et al. (2013). Thymoquinone induces apoptosis in human colon cancer cells through mitochondrial depolarization and caspase activation. International Journal of Oncology, 42(3), 959–965.

40. Peng, L., et al. (2019). Thymoquinone inhibits the proliferation and invasion of ovarian cancer cells by modulating the PI3K/AKT/mTOR pathway. Oncology Reports, 41(1), 281–290.

41. Lei, X., et al. (2020). Thymoquinone inhibits the growth of pancreatic cancer cells by suppressing the Wnt/β-catenin pathway. Cancer Letters, 471, 1–10.

42. Honigberg, L. A., et al. (2010). The Bruton tyrosine kinase inhibitor PCI-32765 blocks B-cell activation and is efficacious in models of autoimmune disease and B-cell malignancy. Proceedings of the National Academy of Sciences, 107(29), 13075–13080.

